# Gasdermin D promotes influenza virus-induced mortality through neutrophil amplification of inflammation

**DOI:** 10.1101/2023.03.08.531787

**Authors:** Samuel Speaks, Ashley Zani, Abigail Solstad, Adam D. Kenney, Matthew I. McFadden, Lizhi Zhang, Adrian C. Eddy, Amal O. Amer, Richard Robinson, Chuanxi Cai, Jianjie Ma, Emily A. Hemann, Adriana Forero, Jacob S. Yount

**Author notes:** Equal Contribution (Samuel Speaks, Ashley Zani).

## Abstract

Influenza virus activates cellular inflammasome pathways, which can be either beneficial or detrimental to infection outcomes. Here, we investigated the role of the inflammasome-activated pore-forming protein gasdermin D (GSDMD) during infection. Ablation of GSDMD in knockout (KO) mice significantly attenuated virus-induced weight loss, lung dysfunction, lung histopathology, and mortality compared with wild type (WT) mice, despite similar viral loads. Infected GSDMD KO mice exhibited decreased inflammatory gene signatures revealed by lung transcriptomics, which also implicated a diminished neutrophil response. Importantly, neutrophil depletion in infected WT mice recapitulated the reduced mortality and lung inflammation observed in GSDMD KO animals, while having no additional protective effects in GSDMD KOs. These findings reveal a new function for GSDMD in promoting lung neutrophil responses that amplify influenza virus-induced inflammation and pathogenesis. Targeting the GSDMD/neutrophil axis may provide a new therapeutic avenue for treating severe influenza.

## Introduction

Influenza A virus (IAV) infection remains a threat to global public health with seasonal epidemics affecting nearly 10% of the world’s population annually and emergent global pandemics remaining an ever-present threat^1, 2^. Exuberant inflammatory immune responses and excessive tissue damage often characterize severe IAV infections. Indeed, IAV is known to trigger multiple inflammatory pathways, including the activation of cellular inflammasomes through several mechanisms^3^. However, whether inflammasome activation is beneficial or detrimental to the outcome of infection is a topic of debate^4–9^, which may indicate that distinct pathways and molecules downstream of inflammasome activation have divergent effects.

The NLRP3 inflammasome is amongst the best characterized inflammasomes in the context of IAV infection and its activation is a double-edged sword, driving both beneficial effects and pathological inflammation^3, 6^. NLRP3 knockout (KO) mice exhibit accelerated death upon IAV infection, a phenotype that is associated with a decrease in protective proinflammatory cytokine secretion^5^. On the other hand, partial blockade of the NLRP3 inflammasome via chemical inhibition at day 7 post infection is beneficial during influenza virus infection^6, 10–12^. These improved outcomes were also associated with decreased production of inflammatory cytokines and chemokines. In what may be a related finding, we have shown dramatically reduced morbidity and mortality in mice treated with an experimental therapeutic protein, recombinant human Mitsugumin 53 (MG53, also known as TRIM72^13^), following otherwise lethal IAV infection. MG53-mediated protection correlated with significantly lower levels of NLRP3 upregulation in infected lungs^14^. Together, these studies suggest that excessive inflammasome activation contributes to pathology and death during IAV infection.

Gasdermin D (GSDMD) is an effector protein that is cleaved by inflammasome-activated caspases, allowing the liberated N-terminal GSDMD fragment to assemble into oligomers that form ring-shaped pores in the plasma membrane^15, 16^. GSDMD pores mediate release of specific pro-inflammatory cytokines^17–19^, which can promote leukocyte recruitment and viral clearance^20, 21^. Unresolved pore formation can lead to release of additional inflammatory cytosolic components and pyroptotic cell death^15, 22–24^. *In vitro*, GSDMD was shown to be non-essential for macrophage death induced by influenza virus infection, though effects on cytokine and chemokine responses were not examined^8^. GSDMD has been subsequently overlooked in influenza virus studies *in vivo* that have utilized genetic knockouts of inflammasome components and effectors. Nonetheless, the potential role for GSDMD in driving inflammation during viral infection *in vivo* warrants further investigation.

In the present study, we explored the role of GSDMD in IAV infections using GSDMD KO mice and discovered that loss of GSDMD decreases lung inflammation, lung dysfunction, and overall mortality during this viral infection, phenotypes that were linked to prevention of excessive neutrophil activation that amplifies lung inflammation. The observed decreases in inflammation in the absence of GSDMD did not affect virus loads or hinder recovery from infection. Thus, our study identifies GSDMD as an exciting potential target for decreasing pathological neutrophil activities and the severity of influenza virus infections.

## Results

### Reduced morbidity and mortality in IAV infected GSDMD KO mice

To investigate the roles of GSDMD during IAV infection, we infected wild type (WT) and GSDMD KO mice with influenza virus strain A/Puerto Rico/8/34 (H1N1) (referred to hereafter as PR8). We first examined lungs for total GSDMD levels, as well as cleaved GSDMD N-terminal fragment indicative of activation, at day 7 post infection via western blotting. Cleaved GSDMD was detected in the lungs of infected WT mice, as expected^14^, whereas only full-length GSDMD could be seen in lungs from mock-infected WT mice (**Fig 1A**). GSDMD could not be detected in lungs from infected GSDMD KO mice, confirming genetic ablation in these animals (**Fig 1A**). GSDMD KO mice lost significantly less weight than their WT counterparts (**Fig 1B**) though the observed weight loss difference underestimates the protective effects of GSDMD deficiency because WT mice had significantly higher mortality rates throughout infection; thus, the sickest animals were removed from the study as they succumbed to infection, skewing the subsequent weight averages. Indeed, over 60% of WT mice succumbed to infection, compared to roughly 10% of GSDMD KO animals, demonstrating a profound protective effect when GSDMD is absent (**Fig 1C**). We next evaluated whether GSDMD impacts viral replication. We found that virus titers in the lungs of WT and KO mice were similar (**Fig 1D**), indicating that GSDMD contributes to the morbidity and mortality of IAV infections without directly affecting virus levels.

**Figure 1:**
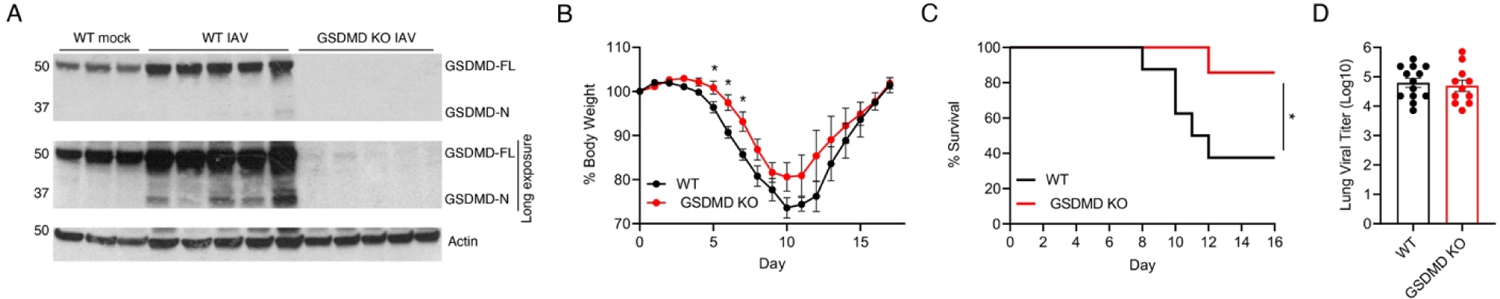
GSDMD promotes inflammation and morbidity following IAV infection. **A-I** WT and GSDMD KO mice were intranasally infected with 50 TCID50 of IAV strain PR8. **A** Western blot of lung lysates taken at day 7 post infection. **B** Weight loss (*p < 0.05 Mann-Whitney test) from 3 independent experiments. **C** Survival curve (*p < 0.05 Log-rank (Mantel-Cox) test) from 3 independent experiments. **D** Viral titers from lung homogenates taken at day 7 post infection from 3 independent experiments.

### GSDMD augments lung pathology during IAV infection

To further explore the protective effects seen in GSDMD KOs during IAV infection, we examined lung sections via hematoxylin and eosin (H&E) staining at day 7 post infection. Although all infected lung sections showed areas of cell infiltration and consolidation, WT lungs exhibited more severe pathology, with thickened alveolar septa, cellular accumulation, and less open airspace as compared to GSDMD KO lungs (**Fig 2A**). Areas of cellular consolidation versus open airspace were quantified to determine relative lung cellularity as a measure of pathology in individual mice ^9, 25^. GSDMD KO mice indeed showed decreased lung pathology via this unbiased quantification method as compared to WT mice (**Fig 2B**).

**Figure 2:**
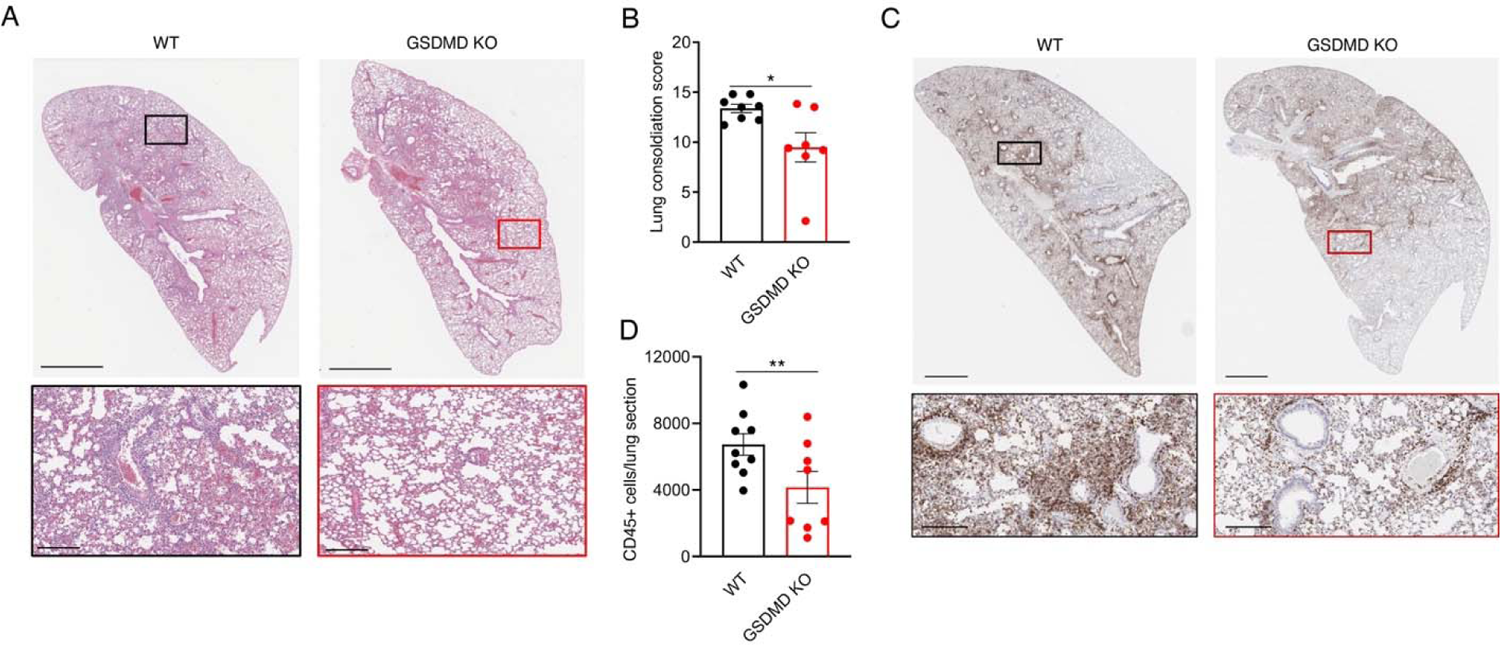
GSDMD exacerbates lung pathology and immune cell infiltration following IAV infection. **A** Representative H&E images for WT and GSDMD KO mice at day 7 post-infection. Insets show magnified areas of cell infiltration. Scale bars represent 2mm and 200um, respectively. **B** Quantification of H&E staining for entire lung sections of multiple animals in two separate experiments (*p < 0.05, t-test). **C** Representative CD45^+^ IHC images for WT and GSDMD KO mice at day 7 post-infection. Insets show magnified areas of CD45^+^ immune cell infiltration. Scale bars represent 2mm and 200um, respectively. **D** Quantification of CD45 staining for multiple animals in two separate experiments (** p < 0.01, t-test).

We next stained for the hematopoietic immune cell marker CD45 in lung sections to assess whether the decreased cellularity and tissue consolidation observed in GSDMD KO lungs was related to dampened immune cell infiltration. We found that infected GSDMD KO mouse lung sections suggested a modest decrease in CD45^+^ immune cells as compared to infected WT lungs (**Fig 2C**). Notably, immune cells were often clustered around the peribronchiole and perivascular spaces in WT mice compared to more diffuse immune cell staining throughout the lung tissue of GSDMD KO mice. Quantification of CD45 staining in individual mouse lung sections confirmed a significant decrease, on average, in CD45^+^ cells for GSDMD KO relative to WT lungs (**Fig 2D**). These results indicate that loss of GSDMD decreases immune-mediated pathology during influenza virus infection, consistent with our observed protection of GSDMD KO mice from influenza virus-induced morbidity and mortality (**Fig 1B,C**).

### GSDMD KO mice have decreased inflammatory transcriptional responses following IAV infection

To further understand the underlying mechanisms contributing to differences in survival and lung damage between WT and GSDMD KO animals, we compared global gene transcriptional responses of WT and GSDMD KO mice following IAV infection. Quantitative measurement of total lung mRNA expression at day 7 post infection showed distinct transcriptional profiles elicited between WT and GSDMD KO animals as revealed by principal components analysis (**Fig 3A**). Statistical analysis of differential gene expression (fold change |3|, p-adj < 0.01) captured 812 genes that were increased in infected WT mice compared to infected GSDMD KO mice, while expression of 447 genes was increased in KOs relative to WT, for a total of 1259 differentially expressed genes (**Fig 3B, Supplementary Table 1**). Hierarchical clustering of gene expression revealed gene clusters that were highly upregulated (yellow cluster) and moderately upregulated (red cluster) in infected WT mice relative to GSDMD KO mice (**Fig 3C**). Gene ontology enrichment revelaed that highly upregulated genes corresponses to biological processes involved in immune cell chemotaxis, granulocyte infiltration, and antiviral responses while modestly expressed genes are involved in cytokine production and inflammatory responses (**Fig 3D**). Genes found to be repressed in WT infected mice relative to GSDMD KO mice (orange cluster), such as *Kcnh2*, *Scn5a*, and *Sgcg*, were found to associate with muscle function. Gene set enrichment analysis followed by network analysis further confirmed that biological processes associated with inflammation and cell-to-cell communication were dampened by the absence of GSDMD in IAV infection (**Fig 3E**).

**Figure 3:**
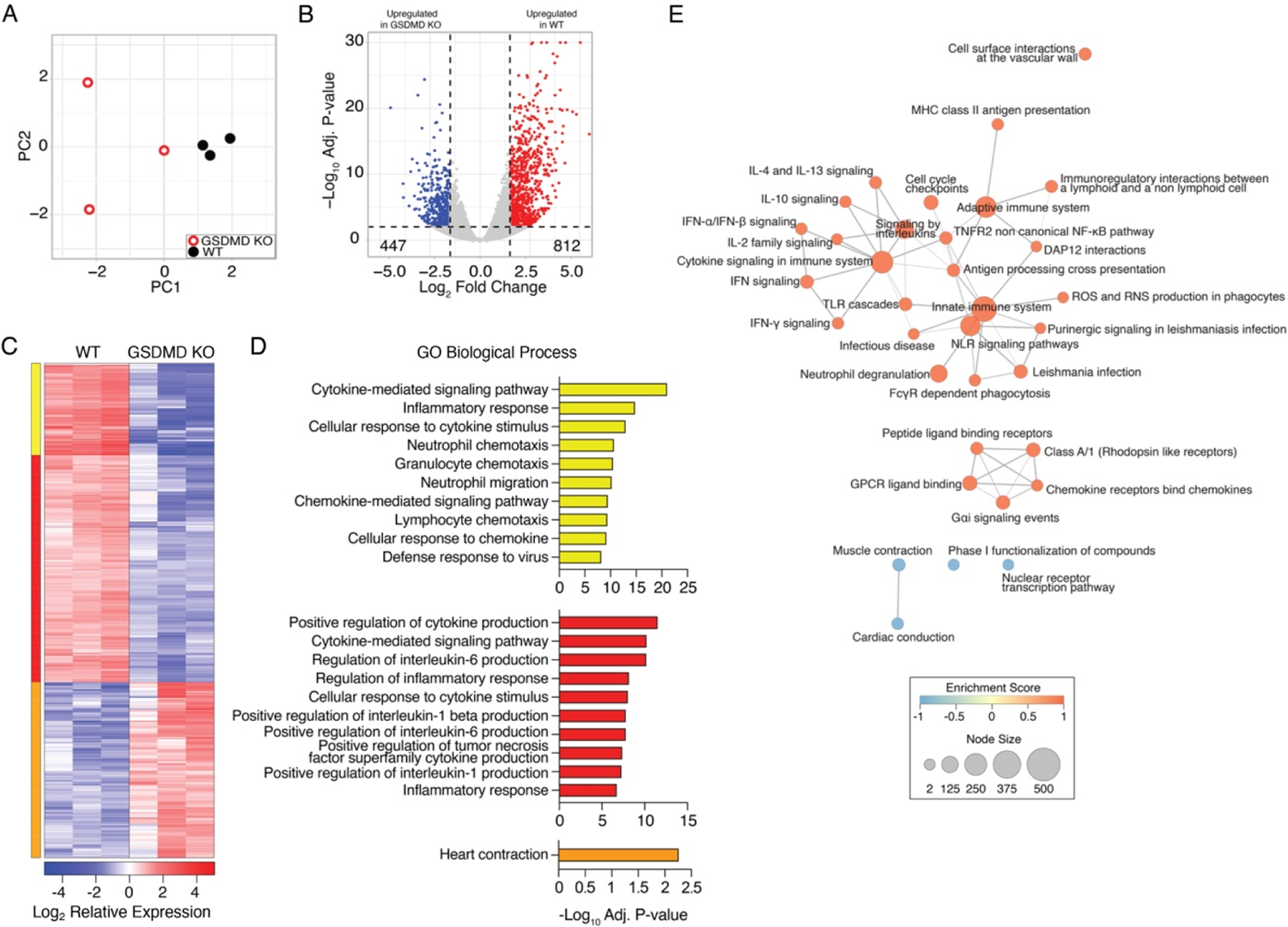
GSDMD promotes inflammatory gene expression programs. WT and GSDMD KO mice were infected with IAV strain PR8 at a dose of 50 TCID50. A-E RNA was extracted from lungs at day 7 post infection and subjected to RNA sequencing. **A** Principal component analysis comparing WT and GSDMD KO lung RNA sequencing results. **B** Volcano plot of differential gene expression (lfc |1.5|, adj p-value <0.01) comparing WT vs GSDMD KOs. Red, 812 genes upregulated in WT vs KO; Blue, 447 genes downregulated in WT vs KO. **C** Hierarchical clustering of differentially expressed genes following IAV infection. Heatmap represents relative gene expression where red indicates genes with upregulated expression and blue indicates genes with downregulated expression. Cluster color indicates genes with similar patterns of expression in WT infected mice compared to GSDMD KO. **D** Gene ontology analysis for genes within each cluster. Bar graphs represent the top 10 enriched GO terms enriched within each cluster as indicated by color. Bar length represent the -log10 adj. p value for significantly enriched pathways (-Log10 adj p-value > 1.3). **E** Network of GO Biological Process terms enriched by gene set enrichment analysis (GSEA). Node size represents number of genes within each pathway. Edges represent the number of shared genes across GO BP terms. Color indicates the GSEA Enrichment Score (ES) where orange indicates positive ES and blue indicates negative ES for transcriptional signatures derived from WT infected lungs relative to GSDMD KO.

### Loss of GSDMD reduces neutrophil activation programs during IAV infection

Additional analysis of RNA sequencing data illustrated that gene expression linked to inflammatory defense responses to virus infection was decreased in GSDMD KO lungs (**Fig 4A**). This gene set included interferons (IFNs), IFN-stimulated chemokines, and classical antiviral IFN-stimulated genes such as MX and OAS family genes (**Fig 4A**). In addition to changes in cytokines and chemokines, functional enrichment analysis of differentially expressed genes that intersect with immune cell specific gene expression (PanglaoDB) suggested significant decreases in myeloid cell associated gene products in GSDMD KO lungs (**Supplementary Fig 1A**). This was consistent with observed enrichment of biological pathways involved in granulocytic neutrophil function in WT versus KO samples as identified by GO Biological Process analysis (**Fig 3D, Fig 4B**), REACTOME analysis, (**Fig 4C, Supplemental Fig 1B**), and Ingenuity Pathway Analysis (**Supplemental Fig 1C)**. These expression differences included genes associated with neutrophil degranulation as well as decreases in genes for chemokines that recruit neutrophils, chemokine receptors expressed on neutrophils, and other molecules associated with neutrophil function (**Fig 4B,C**). Together, these RNA sequencing results broadly identified decreased inflammatory responses during IAV infection in GSDMD KOs and implicated decreased neutrophil activity in mediating the observed differences between WT and KO animals.

**Figure 4:**
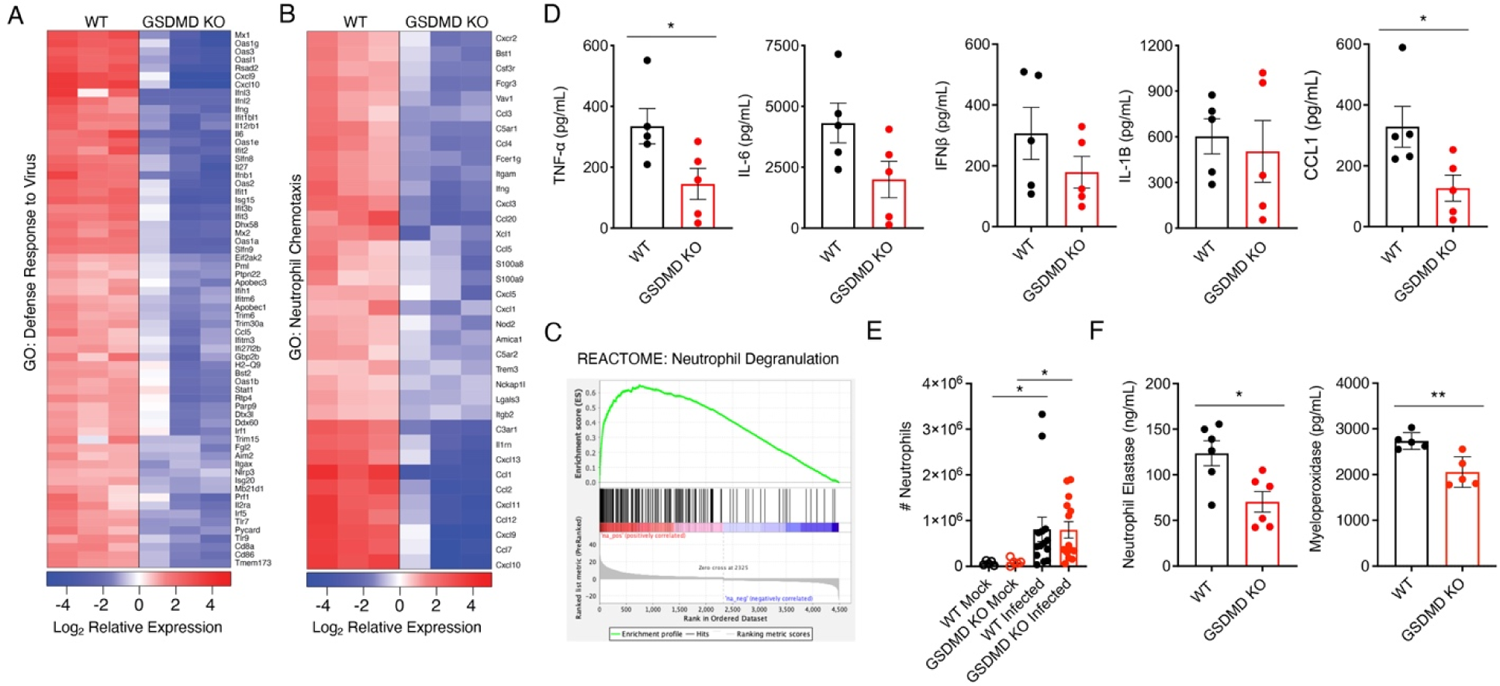
Neutrophil activity is diminished in GSDMD KO mice. **A,B** RNAseq results as in Figure 3 were further analyzed. **A** Heat map for top differentially regulated genes involved with Response to Virus as defined by GO biological process. **B** Heat map for top differentially regulated genes involved with neutrophil chemotaxis as defined by GO biological processes. **C** REACTOME analysis of differentially expressed genes in WT versus GSDMD KO mice identified significant associations with neutrophil degranulation. **D** ELISA quantification of TNFα, IL-6, IFNβ, IL-1β, or CCL1 levels in lung homogenates of WT and GSDMD KO mice at day 7 post infection. (*p < 0.05, t-test). **E** Neutrophil infiltration into the lungs of mock and infected mice quantified by flow cytometry (*p < 0.05, two-way ANOVA and Tukey multiple comparisons test). **F** ELISA quantification of neutrophil elastase levels in lung homogenates of infected WT and GSDMD KO mice at day 7 post infection. (*p < 0.05, **p < 0.01, t-test).

Since our RNAseq analyses revealed a broad attenuation of inflammatory cytokine and chemokine gene programs in GSDMD KO lungs, with CCL1, a chemokine secreted by activated monocytes, macrophages, and T cells, being the most significantly downregulated gene identified (**Fig 4B**), we measured levels of a panel of secreted factors in WT and GSDMD KO lungs during infection. Indeed, GSDMD KO mice had reduced levels of pro-inflammatory cytokines, on average, as measured by ELISA, including IL-6 and IFNβ, with levels of TNFα showing a statistically significant decrease compared to WT mice (**Fig 4D**). Interestingly, levels of IL-1β varied widely between animals and did not correlate with animal genotype (**Fig 4D**) despite the established role of GSDMD in facilitating release of IL-1β^17–19^ and the indication of a decreased IL-1 response in our RNA sequencing results (**Fig 3D**). This is consistent with our previous findings in which GSDMD was required for maximal levels of lung IL-1β early in SARS-CoV-2 infection, but was dispensible later in infection^9^. We also observed that the chemokine CCL1 was significantly decreased in GSDMD KO versus WT lungs (**Fig 4D**), consistent with the major changes observed for this chemokine in our RNA sequencing results (**Fig 3B**). To determine whether human GSDMD functions similarly to its mouse counterpart in promoting cytokine and chemokine secretion during influenza virus infection, we infected WT and GSDMD knockdown (KD) human THP1 macrophages and measured viral protein and cytokine levels after 48 hours. In accord with our *in vivo* results, we saw a decrease in IL-1β, IL-6, TNF, IFNβ and CCL1 levels secreted by GSDMD KD macrophages despite comparable levels of infection as indicated by viral nucleoprotein expression in Western blots (**Supplemental Fig2A-C**). Our data collectively suggest that dampened inflammatory responses in the absence of GSDMD underly the decreased lung pathology and increased survival of GSDMD KO mice following IAV infection.

Next, we examined whether GSDMD was involved in neutrophil recruitment to the lungs during influenza virus infection. We surprisingly found that both WT and GSDMD KO mice exhibited robust neutrophil recruitment to the lungs by day 7 post IAV infection with no statistical difference observed between the genotypes in terms of numbers of neutrophils per lung (**Fig 4E**) or percent of neutrophils relative to total CD45^+^ immune cells (**Supplemental Fig 3**). We also observed similar recruitment of other innate immune cell populations, such as eosinophils, macrophages, or natural killer cells into the lungs, and likewise, recruitment of adaptive CD4 and CD8 T cells and B cells was not impaired in KO animals (**Supplemental Fig 3**). We thus hypothesized that despite similar numbers of neutrophils in the lungs of GSDMD KOs, that perhaps neutrophil functionality was decreased. We then measured levels of neutrophil elastase, a proteolytic enzyme released by activated neutrophils that degrades extracellular matrix and promotess neutrophil extracellular trap formation^26–28^. This indicator of activated neutrophils was significantly decreased in the lungs of infected GSDMD KO as compared to WT tissue (**Fig 4F**). We similarly measured myeloperoxidase, molecule released by azurophilic neutrophil granules that catalyzes production of reactive oxygen intermediates known to contribute to tissue damage, and found that it was also decreased in the absence of GSDMD (**Fig 4F**). In sum, both RNA sequencing results and measurements of neutrophil products indicated decreased neutrophil functionality during infection of GSDMD KO compared to WT lungs.

### Neutrophils increase inflammation and mortality in IAV infection

To probe whether decreased neutrophil activities could explain the decrease in influenza virus infection severity that we observed in the GSDMD KO animals, we depleted neutrophils early in infection of WT mice and measured infection outcomes. Briefly, we treated mice with antibodies targeting neutrophils (α-Ly6G) or isotype control antibody, starting at day 3 post infection through day 8 post infection (**Fig 5A**). This treatment timeline was chosen to allow acute neutrophil responses to proceed uninhibited prior to blockade of a presumptive excessive, pathological response, and also because it represented a test of neutrophil depletion as a therapeutic strategy. We confirmed that antibody-mediated depletion led to a marked decrease in total neutrophil numbers, as well as relative percentage of neutrophils compared to the entire CD45^+^ immune cell population, in the lungs of α-Ly6G treated mice at day 5 post infection (**Fig 5B, Supplementary Fig 4A,B**). The specifity of neutrophil depletion was further confirmed by analyzing total counts and percentages of eosinophils and alveolar macrophages, which were not significantly different between the two treatment groups. (**Supplementary Fig 4A,B**). When examining morbidity between the cohorts of mice, the α-Ly6G treated mice experienced significantly less influenza virus-induced weight loss than the isotype control treated group (**Fig 5C**). Remarkably, neutrophil depletion was completely protective against lethal infection, whereas we observed 60% lethality in isotype control-treated mice (**Fig 5D**), a result that mirrored the protective effects seen in GSDMD KO mice (**Fig 1C**).

**Figure 5:**
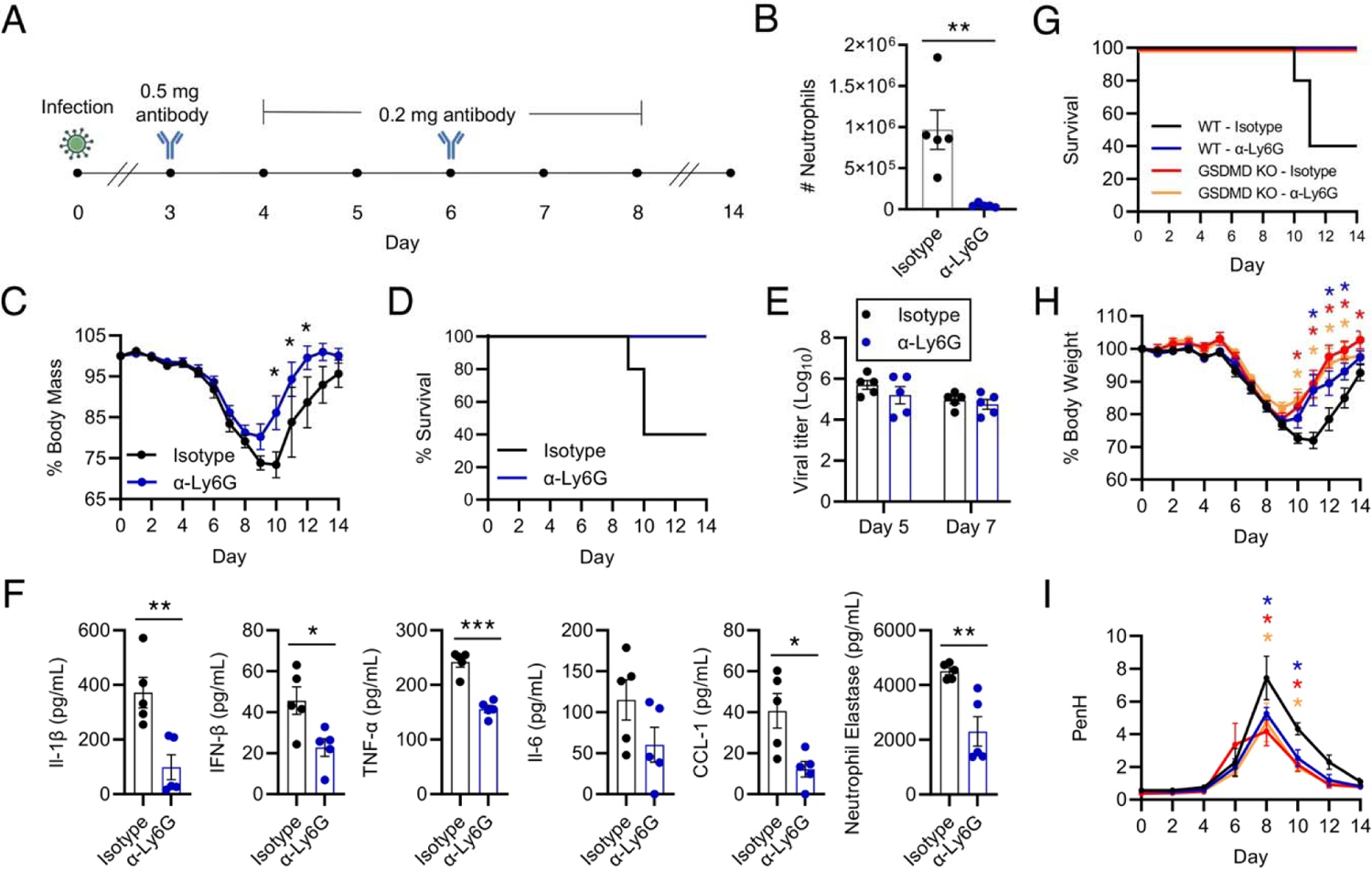
Neutrophils potentiate severe influenza virus-associated illness. **A** WT mice were infected with 50 TCID50 of PR8. Mice were treated via intraperitoneal (IP) injection with 0.5mg of α-Ly6G antibody or isotype control antibody on day three post infection, followed by 0.2mg of the corresponding antibody on days four through eight. **B** Neutrophil number in the lungs of WT and GSDMD KO mice on day 5 post infection as in **A**, quantified by flow cytometry. (**p < 0.005, t-test) **C** Weight loss of mice infected as in **A** that were followed until recovery (*p < 0.005, Mann-Whitney test). **D** Survival analysis of mice as infected in **A**. **E** Viral titers from lung homogenates at day 5 and day 7 post infection. **F** ELISAs of pro-inflammatory cytokines performed on lung homogenates from day 7 post infection (*p < 0.05, ** p < 0.01, *** p < 0.001, t-test). **G** Survival analysis of infected WT and GSDMD KO mice treated with α-Ly6G antibody or isotype control antibody as in **A**. **H** Weight loss of mice infected as in **G** that were followed until recovery (*p < 0.05, Mann-Whitney test, colored asterisks represent comparisons to WT - Isotype). Colored lines correspond to groups as in **G)**. **I** PenH, an indicator of airway resistance, in WT and GSDMD KO mice as infected in **G** measured by whole body plethysmography (*p < 0.05, two-way ANOVA and Tukey multiple comparisons test, colored asterisks represent comparisons to WT - Isotype). Colored lines in **H** and **I** correspond to groups as in **G**.

Also similar to results comparing GSDMD KO and WT mice mice, viral titers in the lungs of neutrophil-depleted and isotype control-treated mice were not statistically different from one another at either day 5 or day 7 post infection (**Fig 5E**). We further observed that levels of inflammatory cytokines, including IL-1β, IL-6, IFNβ and TNF-α, were significantly decreased in the lungs of neutrophil-depleted mice compared to controls (**Fig 5F**). Likewise, CCL1 levels were commensurately decreased following α-Ly6G treatment (**Fig 5F**). These results suggest that neutrophils overall potentiate a postive-feedback loop of inflammation in IAV infected lungs. Finally, we quantified neutrophil elastase in the lungs of α-Ly6G treated and isotype control treated mice to further confirm that neutrophil elastase serves as a marker of neutrophil presence and activity. Indeed, we saw decreased neutrophil elastase in the lungs of α-Ly6G treated mice (**Fig 5F)**, reinforcing our data identifying a similar change in GSDMD KO versus WT mice (**Fig 4F).**

To gain further insight into whether the protection provided by loss of GSDMD during IAV infection was indeed afforded by a decrease in neutrophils, we depleted neutrophils from both WT and GSDMD KO animals during IAV infection (**Supplementary Fig 5**). We again observed that neutrophil depletion or loss of GSDMD protected mice from death and severe weight loss (**Fig 5G,H**). Neutrophil-depleted GSDMD KO mice similarly survived infection but did not show an added benefit over GSDMD KO alone in terms of weight loss (**Fig 5G,H**). We also measured indicators of lung function via full body plethysmography of these mice. PenH, a surrogate measure of airway resistance, increased throughout infection of WT isotype control-treated mice, peaking on day 8 post infection (**Fig 5I**). WT mice with neutrophil depletion, GSDMD KO control mice, and GSDMD KO mice with neutrophil depletion each showed less decline in lung function allowing a more rapid recovery to baseline lung function compared to WT control-treated mice (**Fig 5I**). These plethysmography measurements 1) corroborate our lung histopathology data (**Fig 2**), 2) demonstrate that loss of GSDMD preserves lung function during IAV infection, 3) demonstrate that depletion of neutrophils also benefits lung function during IAV infection, and 4) support the notion that inflammatory, lung damaging effects of GSDMD during IAV are mediated largely by neutrophils. In sum, our results demonstrate that neutrophil recruitment to the lungs during influenza virus infection is largely dependent on GSDMD and that neutrophils amplify the inflammatory response and lung dysfunction during IAV infection.

## Discussion

We discovered that the inflammasome effector, GSDMD, has a detrimental impact on outcomes of influenza virus infection. In the absence of GSDMD, mice experienced reduced morbidity and mortality despite viral burden in the lungs being unaffected (**Fig 1**). Indeed, GSDMD profoundly contributed to lung inflammation, neutrophil activity, lung dysfunction, and histopathology (**Fig 1-5**). The improved outcome of GSDMD KO mice following influenza virus infection compared to WT mice corresponded to global transcriptional decreases in a multitude of tissue-damaging inflammatory pathways (**Fig 3,4**). Additionally, impaired neutrophil activity gene expression programs in the absence of GSDMD correlated with improved disease outcome, which we subsequently substantiated via exogenous depletion of these immune cells (**Fig 5**). Of the numerous immune cell types important for control of influenza virus infection, neutrophils are among the first to infiltrate sites of infection^29^. Beneficial functions of neutrophils include phagocytic activity, cytokine and chemokine release, and formation of neutrophil extracellular traps, which are traditionally associated with antibacterial activities but also promote an antiviral state^30–34^. Conversely, other studies have highlighted the potentially detrimental effect of unchecked neutrophil recruitment characterized by hyperinflammation and lung damage^4, 35–38^. Our data add to a growing number of studies implicating excessive neutrophil activities as a driver of poor disease outcomes^35–37^.

Importantly, GSDMD was also required for maximal pro-inflammatory cytokine secretion by human macrophages infected with influenza virus, suggesting that GSDMD likely also promotes inflammation in human influenza virus infections (**Supplemental Figure 2**). This paralells recent data showing GSDMD co-localization with pyroptotic macrophages in the lungs of severely infected macaques, which also exhibited widespread expression of pro-inflammatory cytokines^4^. In addition to providing important parallels between murine, non-human primate, and human GSDMD, our human macrophage data suggest that GSDMD regulates cytokine production/secretion in macrophages, which are known targets of influenza virus infections^39, 40^.

Influenza virus infection is among the most common causes of acute respiratory distress syndrome (ARDS) in adults^41^. Development of ARDS is marked by bilateral edema and worsened oxygen delivery^42^, and a major contributor to these criteria is the unchecked inflammation caused by cellular immune responses activated to control replicating virus. Specifically, neutrophil infiltration and abundance is correlated with the severity of ARDS^43^, and neutrophil secretion of tissue damaging enzymes, such as neutrophil elastase and myeloperoxidase, and inflammatory extracellular traps can exacerbate progression of ARDS^41^. Likewise, NETosis promoted by GSDMD was recently identified as a driver of disease progression in an LPS-induced model of ARDS^44^. Given that GSDMD is a broad promoter of inflammation in response to infection and that there are numerous infectious causes of ARDS, our results suggest that the GSDMD/neutrophil axis may be a potential therapeutic target for mitigating ARDS progression in the context of other respiratory infections in addition to influenza virus.

While GSDMD is the final effector protein in the pyroptosis pathway initiated by inflammasome activation, other intermediate proteins are likely contributing to the outcome of IAV infection. Indeed, Caspase-1/11 KO mice experience exacerbated acute illness and reduced adaptive immunity to influenza virus^45, 46^. The requirement for specific caspases in activation of lung GSDMD during IAV infection remains to be investigated. Interestingly, neutrophil elastase has also been demonstrated to be capable of cleaving and activating GSDMD^28, 47^, which may explain the opposing outcomes of Caspase-1/11 KO versus GSDMD KO in IAV infections.

Unexpectedly, IL-1β levels were unaffected by the absence of GSDMD, despite its canonical role in IL-1β release^17–19^, suggesting that there there are redundant mechanisms for pyroptotic cytokine release during IAV infection, at least at late timepoints following infection. Indeed, these results are in accord with our published findings that IL-1β levels in the lung at day 2 post infection with SARS-CoV-2 are dependent on GSDMD, but that IL-1β release occurs in a GSDMD-independent manner at later times post infection^9^. Interestingly, we found that loss of GSDMD had no significant impact on SARS-CoV-2 disease severity^9^, highlighting intriguing differences in the pathological mechanisms activated by SARS-CoV-2 versus IAV that warrant further study. In addition, we discovered that GSDMD promotes IFNβ expression and production (**Fig 3, Supplementary Fig 2C)**, further suggesting that this effector protein potentiates pro-inflammatory responses beyond release of canonical inflammasome molecules. Attenuation of several pro-inflammatory cytokine pathways, including IFNβ, in GSDMD KO mice had no appreciable effect on virus replication. However, it is well established that the pathological, tissue-damaging effects of IFNs continue to accumulate beyond the point at which their antiviral effects peak^48–50^. These data indicate that loss of GSDMD during IAV infection balances the inflammatory response toward a less tissue-damaging phenotype while maintaining a response sufficient to combat virus replication.

Overall, we discovered that GSDMD promotes inflammatory lung pathology during influenza virus infection. Importantly, GSDMD is dispensible for virus control and recovery from infection, but exacerbates severe illness, thus identifying it as a promising candidate for host-directed therapeutic targeting during IAV infection.

## Methods

### THP-1 Cell culture

Vector control and and GSDMD KD THP-1 cells were generated by lentiviral shRNA-mediated targeting and were generously provided by Dr. Amal Amer at The Ohio State University. Cells were maintained in RPMI 1640 medium (Fisher scientific) supplemented with 10% EquaFETAL bovine serum (Atlas Biologicals). All cells were cultured in a humidified incubator at 37 °C with 5% CO2. Cells were additionally treated with 25 nM phorbol myristate acetate (PMA) for 3 days as previously described ^51^ to allow for differentiation into macrophages.

### Virus stocks and in vitro infection

Influenza virus A/Puerto Rico/8/34 (H1N1) (PR8, provided by Dr. Thomas Moran of the Icahn School of Medicine at Mt. Sinai) was propagated in embryonated chicken eggs (Charles River Laboratories) and titered as described previously ^52^. Infection of THP-1 cells with PR8 was done at an MOI of 10 for 48hrs.

### Mouse studies

WT and GSDMD KO mice were purchased from Jackson Laboratories. Mice were infected intranasally under anesthesia with isoflurane according to protocols approved by the Ohio State University Institutional Animal Care and Use Committee. Mouse infections were performed with a dose of 50 TCID50 of PR8. For neutrophil depletion experiments, WT mice were treated with 0.5 mg anti-mouse Ly6G antibody (InVivoMAb, BE0075-1) or 0.5 mg rat IgG2a isotype control, anti-trinitrophenol antibody (InVivoMAb, BE0089) via intraperitoneal injection on day 3 post infection. From days 4 through 8 post infection, mice were injected with 0.2 mg of the same antibodies. For determining lung titers and cytokine levels, tissues were collected and homogenized in 1ml of PBS, flash-frozen, and stored at −80°C prior to titering on MDCK cells or analysis via ELISA. TCID50 values were calculated using the Reed-Muench method.

### Western blotting

For detection of protein expression in lung tissue, samples were lysed in a 1% SDS buffer (1% SDS, 50mM triethanolamine pH 7.4, 150mM NaCl) containing a cocktail of phosphatase protease inhibitors (Sigma, 4906845001) and protease inhibitors (Thermo Scientific, A32965). The lysates were centrifuged at 15,000 rpm for 10 min and soluble protein supernatents were used for Western blot analysis. Equal amounts of protein (30ug) were separated by SDS-PAGE and transferred onto membranes. Membranes were blocked with 10% non-fat milk in Phosphate-buffered saline with 0.1% Tween-20 (PBST) and probed with antibodies against GSDMD (abcam, ab219800) and actin (abcam, ab3280). For Western blotting of THP-1 cells, samples were lysed in 1% SDS containing protease inhibitors prior to SDS-PAGE separation as described above. Membranes were probed with antibodies against influenza virus nucleoprotein (abcam, ab20343), GSDMD (abcam, ab210070), and GAPDH (Thermo Scientific, ZG003).

### Single and multiplex ELISAs

human IL-1β, IL-6, CCL1, and TNF, and mouse IFNβ and CCL1 ELISAs were performed on supernatent from lung homogenates or cell culture supernatants using the respective R&D Systems Duoset ELISA kits (catalogue # DY201, DY206, DY8234, DY272, and DY845) according to manufacturer’s instructions. Mouse TNF-α, IL-6, and IL-1β levels were also assayed via multiplex ELISA (Mesoscale Diagnostics, mouse V-plex pro-inflammatory panel 1, K15048D) performed by the OSUMC Center for Clinical Research Management core facility.

### Flow cytometry

On day 7 post influenza virus infection, mice were euthanized and lungs were homogenized in GentleMACS C tubes (Miltenyi) containing Dnase I (Sigma Aldrich) and type IV collagenase (Worthington Biochemical Corporation). Lungs were processed into single cell suspensions using GentleMACS equipment. Single cell homogenates were blocked in CellBlox (Invitrogen) and viability stain was performed using eFluor 780 Fixable Viability Dye (Invitrogen) according to the manufacturer’s instructions. Cells were washed and stained with Invitrogen antibodies for CD45, Ly6G and Ly6C (catalogue number M001T02R02, 46-9668-82, and 48-5932-82, respectively) for 30 minutes on ice. Cells were fixed with BD FACS lysis buffer for 10 minutes at 4°C, resuspended in PBS, and run on a Cytek Aurora instrument. Count beads (Invitrogen) were added to each sample for quantification of total cell number. Samples were analyzed using FlowJo Software (10.8.1).

### Histology

For lung histology (H&E and anti-CD45 staining), lung tissue samples were fixed in 10% neutral-buffered formalin at 4°C for 24 hours and then transferred to 70% ethanol. Lungs were embedded in paraffin, sectioned, stained, and imaged by Histowiz (Histowiz.com, Brooklyn, NY, USA). Unbiased electronic quantification of resultant images was performed using ImageJ software and the color deconvolution method as described previously ^9, 25, 53, 54^.

### RNA sequencing and Functional Enrichment Analysis

RNA was extracted from lung tissue using TRIzol (Invitrogen). RNA library preparation and sequencing was performed by GENEWIZ (GENEWIZ LLC./Azenta US, Inc South Plainfield, NJ, USA) with the following methods, as provided by GENEWIZ: “The RNA samples received were quantified using Qubit 2.0 Fluorometer (ThermoFisher Scientific, Waltham, MA, USA) and RNA integrity was checked using TapeStation (Agilent Technologies, Palo Alto, CA, USA). The RNA sequencing libraries were prepared using the NEBNext Ultra II RNA Library Prep Kit for Illumina using manufacturer’s instructions (New England Biolabs, Ipswich, MA, USA). Briefly, mRNAs were initially enriched with Oligod(T) beads. Enriched mRNAs were fragmented for 15 minutes at 94°C. First strand and second strand cDNA were subsequently synthesized. cDNA fragments were end repaired and adenylated at 3’ends, and universal adapters were ligated to cDNA fragments, followed by index addition and library enrichment by PCR with limited cycles. The sequencing libraries were validated on the Agilent TapeStation (Agilent Technologies, Palo Alto, CA, USA), and quantified by using Qubit 2.0 Fluorometer (ThermoFisher Scientific, Waltham, MA, USA) as well as by quantitative PCR (KAPA Biosystems, Wilmington, MA, USA). The sequencing libraries were multiplexed and clustered onto a flowcell. After clustering, the flowcell was loaded onto the Illumina HiSeq instrument according to manufacturer’s instructions. The samples were sequenced using a 2×150bp Paired End (PE) configuration. Image analysis and base calling were conducted by the HiSeq Control Software (HCS). Raw sequence data (.bcl files) generated from Illumina HiSeq was converted into fastq files and de-multiplexed using Illumina bcl2fastq 2.17 software. One mis-match was allowed for index sequence identification.” Raw fastq files were processed, aligned and quantified with a HyperScale architecture developed by ROSALIND (ROSALIND, Inc. San Diego, CA, https://rosalind.bio/). Read Distribution percentages, violin plots, identity heatmaps, and sample MDS plots were generated as part of the quality control step. Statistical analysis for differential gene expression was performed using the “limma” R library ^55^. The principal component analysis, volcano plots, and heatmaps were formatted in R using numerical values provided by ROSALIND. The topGO R library was used to determine local similarities and dependencies between GO terms in order to perform Elim pruning correction. Additional analysis was performed using the REACTOME database ^56^, PanglaoDB ^57^, and Ingenuity Pathway Analysis (Qiagen).

## Author Contributions

Conceptualization: AOA, CC, JM, JSY; Data Curation: AF, JSY; Formal analysis: SS, AZ, AS, ADK, RR, EAH, AF, JSY; Investigation: SS, AZ, AS, ADK, LZ, ACE, JK, EAH; Visualization: SS, AZ, AS, ADK, AF, JSY; Writing – Original Draft: SS, AZ, ADK, JM, AF, JSY; Supervision: JSY; Project Administration: JSY; Funding Acquisition: JSY; Resources: AOA, CC, JM.

## Acknowledgments

Research in the Yount laboratory is supported by NIH Grants AI130110, AI151230, HL154001, HL157215, and an American Lung Association COVID-19 and Emerging Respiratory Viruses Research Award. AZ was supported by an NSF-GRFP fellowship grant. ADK is supported by an institutional T32 postdoctoral fellowship (NIH grant AI165391).

---

**Supplementary Figure 1:**
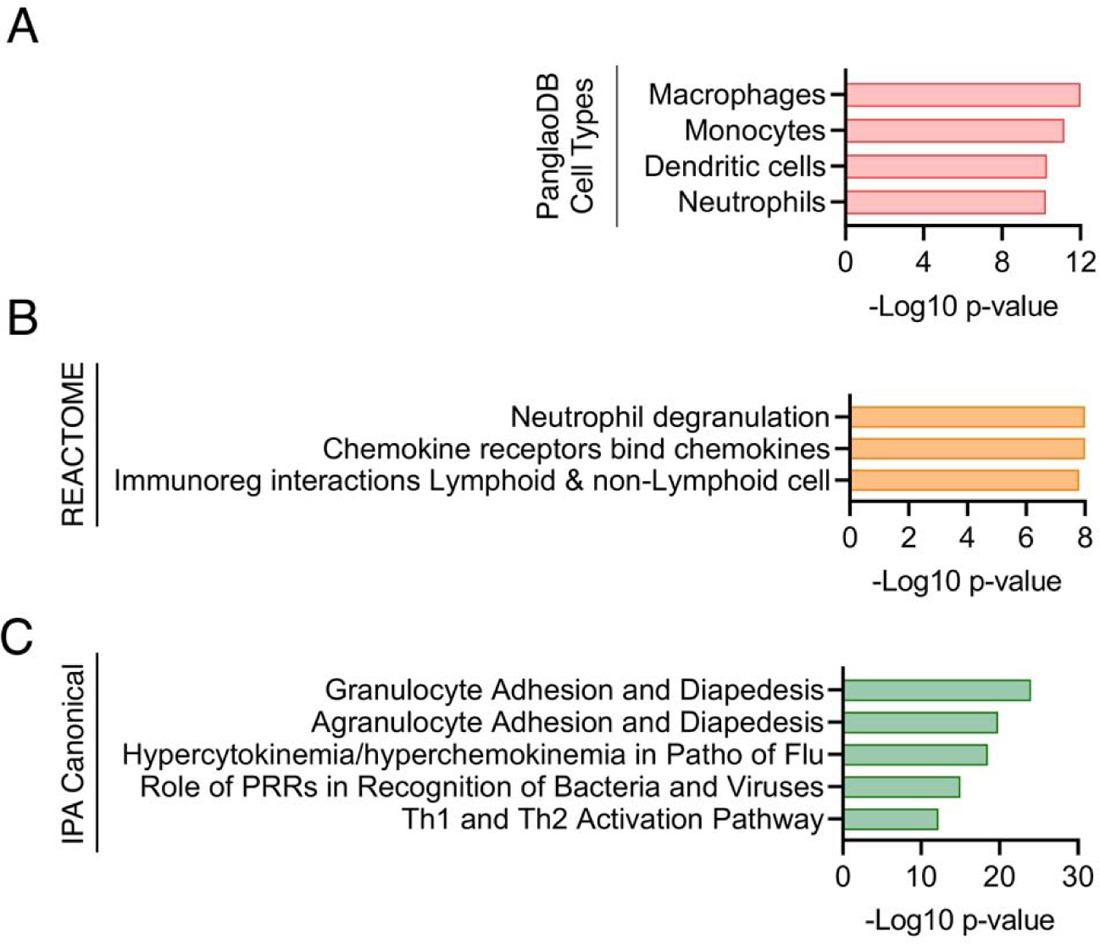
Pathway analysis of gene signatures downregulated in GSDMD KO mice. **A-C** Differentially expressed genes in day 7 post infection GSDMD KO lungs versus WT were subjected to pathway and cell type analysis. **A** Downregualted genes in GSDMD KO versus WT lungs were subjected to PanglaoDB analysis and all significant associations with specific cell types are shown. **B** Significant REACTOME gene set enrichments (p<0.05) for all downregulated genes in GSDMD KO versus WT lungs. **C** All differentially expressed genes were examined using Ingenuity Pathway Analysis and the top five most significant Canonical Pathways are shown.

**Supplemental Figure 2:**
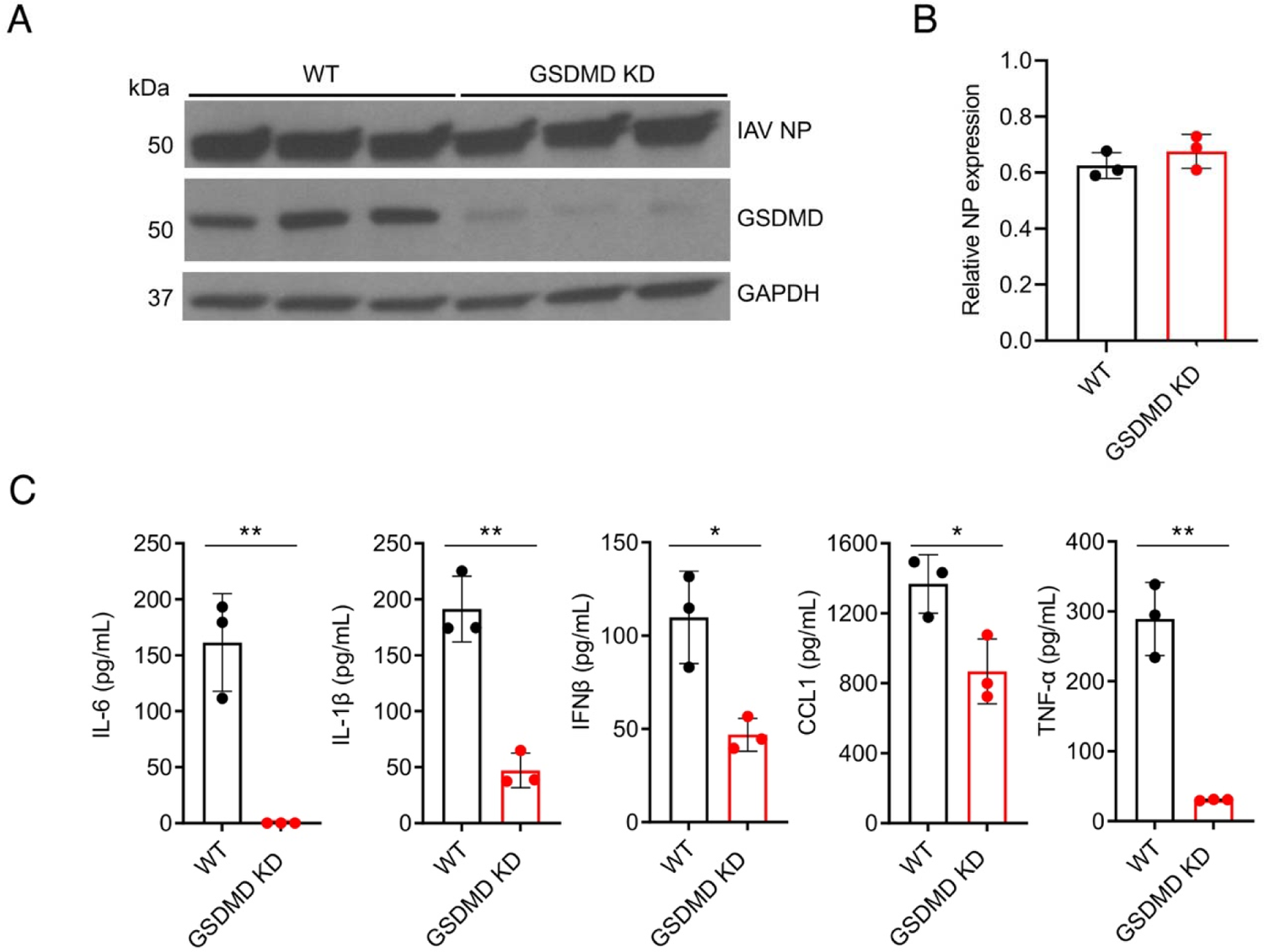
Human GSDMD promotes secretion of pro-inflammatory cytokines *in vitro*. **A** Western blotting for PMA-differentiated WT and GSDMD knockdown (KD) human THP1 macrophages 48 hours post infection with PR8 at an MOI of 10 (NP = influenza virus nucleoprotein). Three technical replicates from one experiment were probed. **B** Densitometry quantifaction of NP levels relative to GAPDH in **A**. **C** ELISA quantification of IL-6, IFNβ, IL-1β, CCL1, or TNF-α levels in supernatants from cells infected as in **A** (*p < 0.05, **p < 0.01, t-test).

**Supplementary Figure 3:**
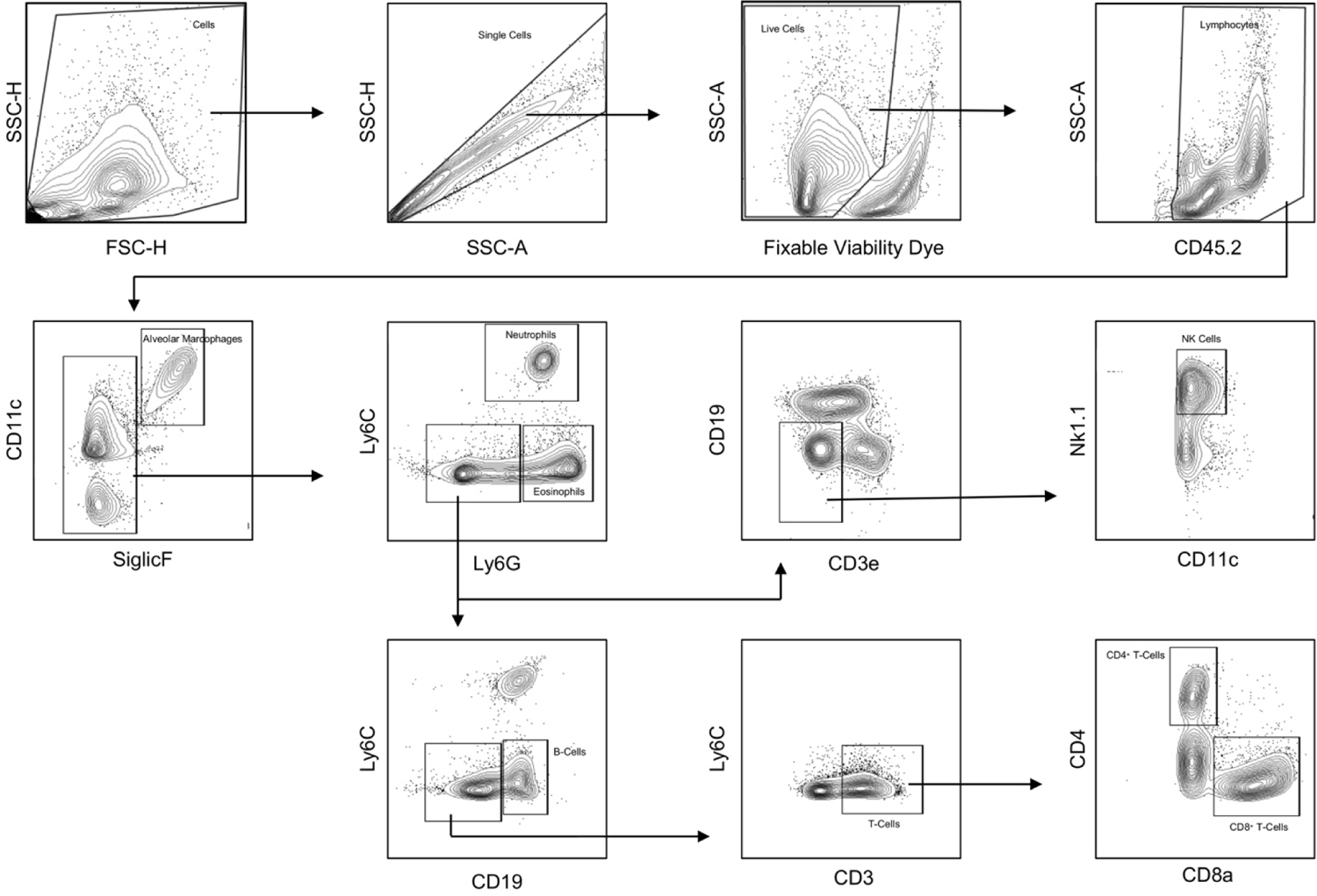
Gating strategy for quantifying immune cell recruitment to mouse lungs during IAV infection.

**Supplementary Figure 4:**
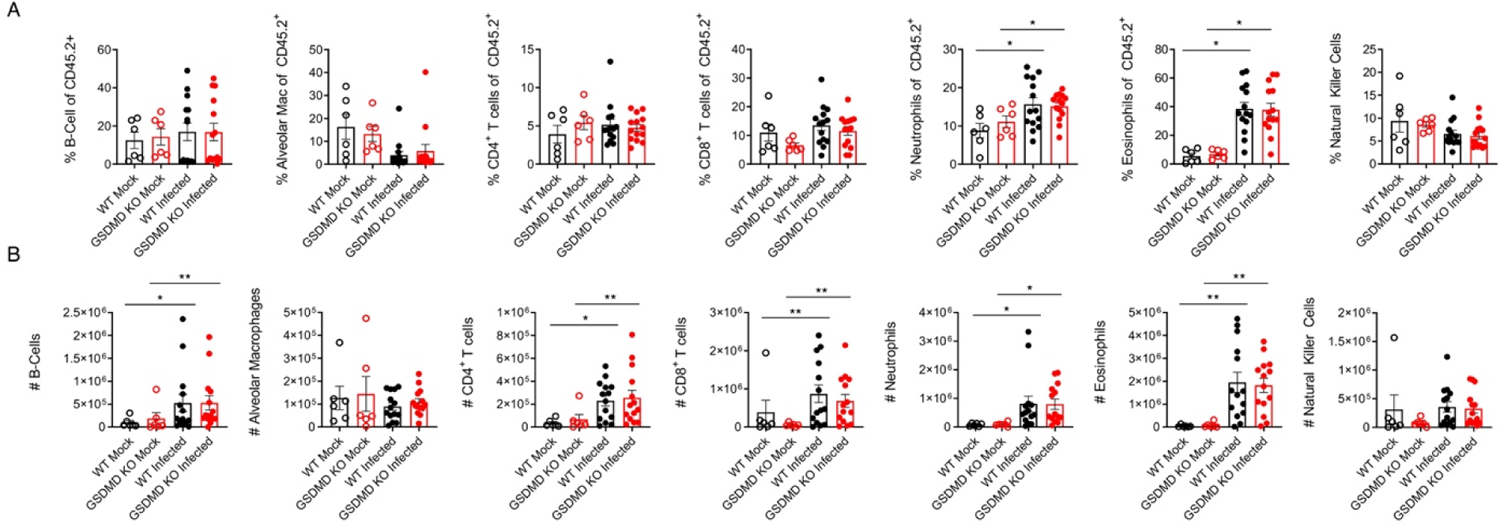
GSDMD does not affect immune cell recruitment to the lung during IAV infection. **A-B** WT and GSDMD KO mice were infected with 50 TCID50 of PR8. Analysis was done on day 7 post infection using the flow cytometry gating strategy show in Supplementary Figure 3. **A** Percentage of indicated cell type relative to all CD45.2^+^ immune cells (*p < 0.05, two-way ANOVA and Tukey multiple comparisons test). **B** Number (#) of infiltrating immune cells into the lungs of mock and infected mice (*p < 0.05, **p < 0.01, two-way ANOVA and Tukey multiple comparisons test). Neutrophil # data is repeated from Fig 4E in main text.

**Supplementary Figure 5:**
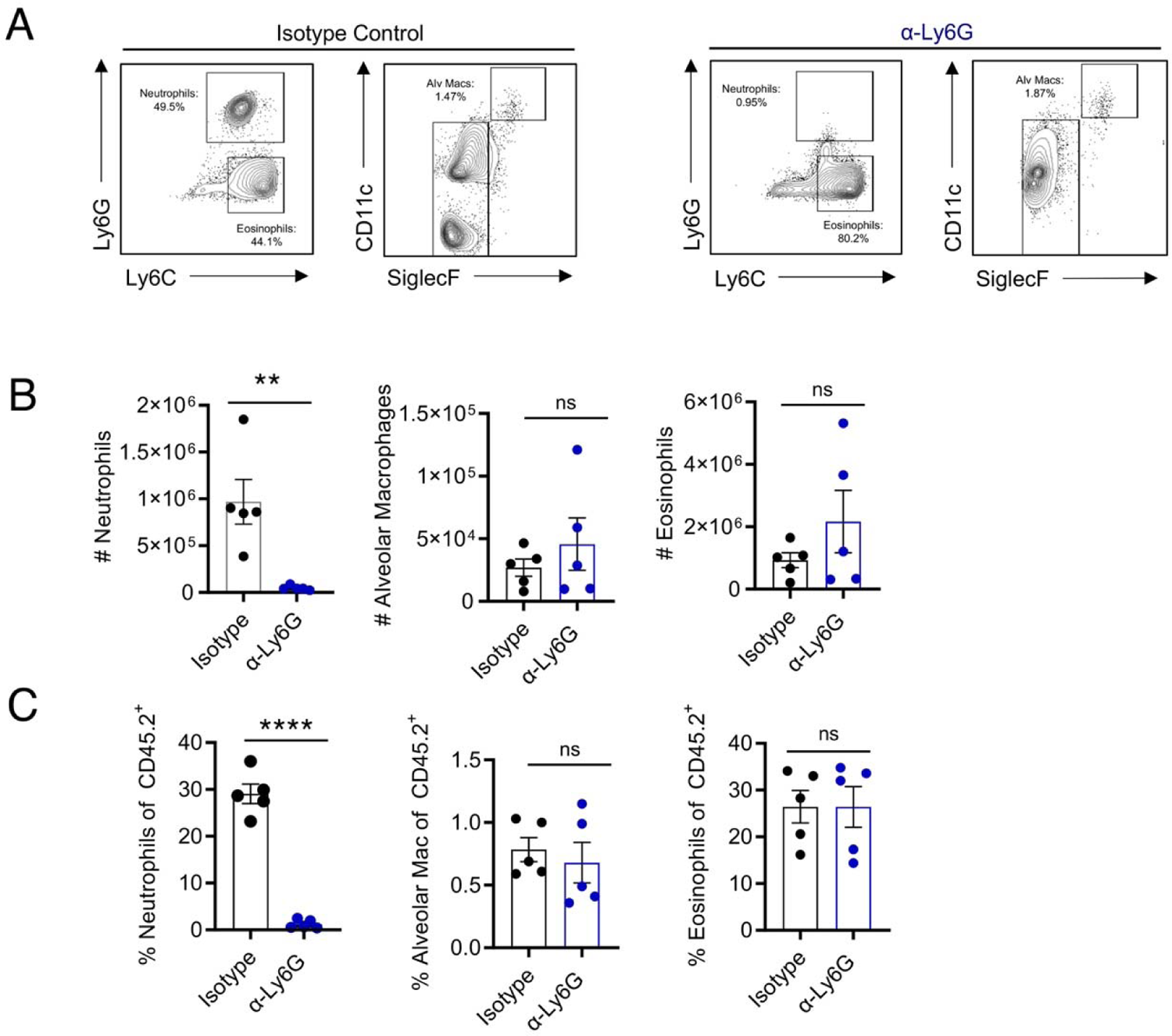
Ly6G antibody treatment depletes neutrophils but not other innate immune cells. **A-C** WT and GSDMD KO mice as infected in Fig 5A. **A** Representative flow cytometryt plots showing neutrophil, eosinophil, and alveolar macrophage gating rom lungs on day 5 post infection. Upstream gating performed as in Supplementary Figure 3. **B** Neutrophil, eosinophil, and alveolar macrophage total number (#) in the lung on day 5 post infection (**p < 0.01, ns, not significant, t-test). Neutrophil # data is repeated from Fig 5B in main text. **C** Neutrophil, eosinophil, and alveolar macrophage percentage of all CD45.2^+^ cells in the lung on day 5 post infection (****p < 0.0001, ns, not significant, t-test).

**Supplementary Figure 6:**
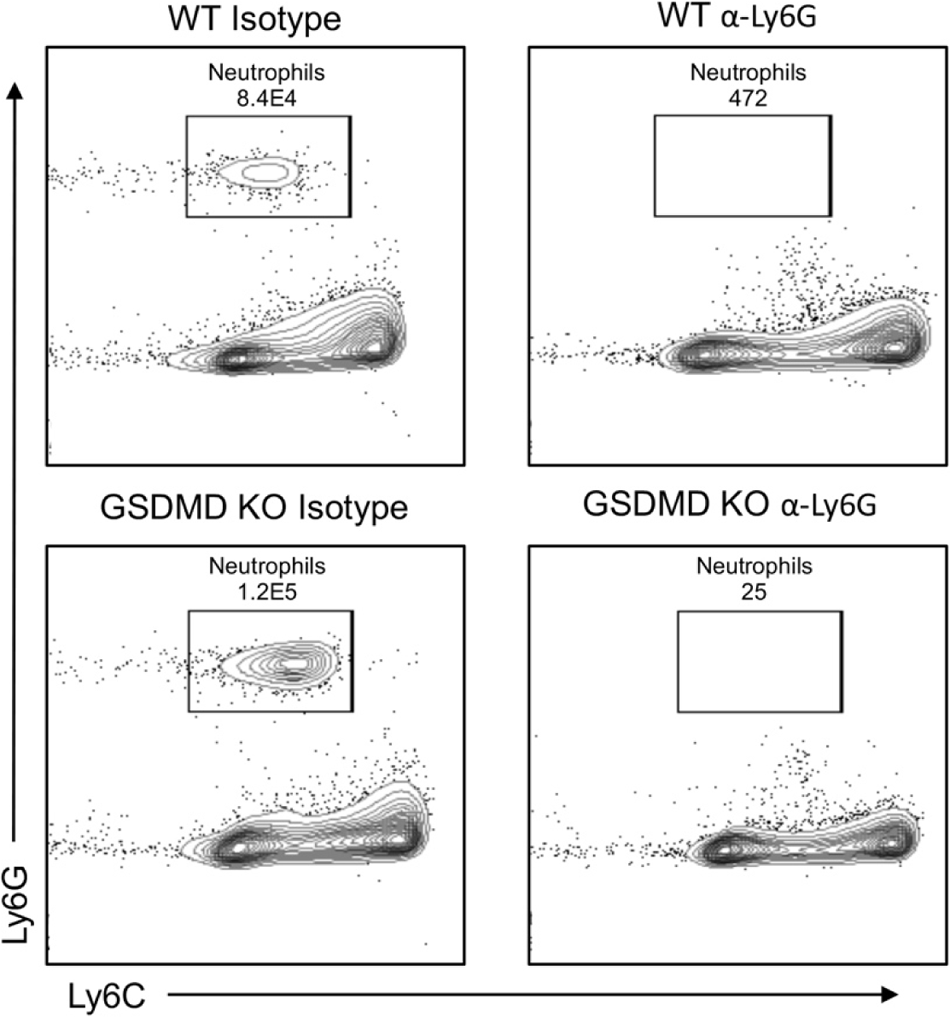
Ly6G antibody treatment depletes neutrophils in GSDMD KO mice as epected. Representative flow cytometry dot plots showing neutrophil gating of single-cell suspensions from the lungs of mice on day 5 post infection as in Fig 5G. These plots represent data from mice randomly chosen to be sacrificed to confirm neutrophil depletion.

